# Regulation of balanced root fungal community of *Vanda falcata* (Orchidaceae) by partitioning its mycorrhizal fungi and ascomycetous fungi across growth and development

**DOI:** 10.1101/2022.01.18.476577

**Authors:** Galih Chersy Pujasatria, Ikuo Nishiguchi, Chihiro Miura, Masahide Yamato, Hironori Kaminaka

## Abstract

Epiphytic orchids are commonly found in exposed environments, which plausibly lead to different root fungal community structures from terrestrial orchids. Until recently, no studies have been conducted to show the fungal community structure during the growth of a photosynthetic and epiphytic orchid in its natural growing state. In this study, the *Vanda falcata* (commonly known as *Neofinetia falcata*), one of Japan’s ornamental orchids, was used to characterize the fungal community structure at different developmental stages. Amplicon sequencing analysis showed that all development stages contain a similar fungal community: Ascomycota dominates half of the community while one-third of the community is Basidiomycota. *Rhizoctonia-*like fungi, a term to group the most common basidiomycetous fungi that form orchid mycorrhiza, exist even in a smaller portion (around a quarter) compared to other Basidiomycota members. While ascomycetous fungi exhibit pathogenicity, two *Ceratobasidium* strains isolated from young and adult plants could initiate seed germination *in vitro*. It was also found that the colonization of mycorrhizal fungi was concentrated in the lower part of the root where it directly attaches to the phorophyte bark, while ascomycetous fungi were distributed in the velamen but never colonized cortical cells. Additionally, lower root parts attached to the bark have denser exodermal passage cells, and these cells were colonized only by mycorrhizal fungi that further infiltrated the cortical area. Therefore, we propose that physical regulation of fungal entry to partition the ascomycetes and mycorrhizal fungi results in the balanced mycorrhizal symbiosis in this orchid.

## Introduction

Orchidaceae, one of the biggest flowering plant families, is widely distributed in almost all parts of the world. However, many orchid species are listed as endangered owing to habitat degradation and dependence on other organisms, i.e., pollinators (Suetsugu et al. 2015; Freitas et al. 2020). Moreover, establishing orchids in the natural habitat is always complicated, even in suitable conditions (Pujasatria et al. 2020). Due to its unique, one-of-a-kind traits, orchid seed germination is often hard to occur naturally. Despite extremely high fecundity, endosperm in orchids’ seeds is almost absent. Consequently, orchids’ seeds are nearly impossible to germinate without nutritional support. For seeds to successfully germinate and grow into mature plants, they must establish orchid mycorrhizal (OM) symbiosis (Arditti and Ghani 2000; Barthlott et al. 2014; Yeh et al. 2019; Pujasatria et al. 2020). Colonization of orchid mycorrhizal fungi (OMF) starts at seed germination and may last until adulthood. As with other mycorrhizae, OMF forms coiled hyphae in its host cells. This structure is termed peloton, where the exchange of nutrients (e.g., carbon, nitrogen, and phosphorus) occurs (Yeh et al. 2019). However, OM is unique because the host eventually digests the pelotons for further nutrient acquisition (Kuga et al. 2014). Based on this, orchids are considered parasitizing the fungus. This behavior, termed mycoheterotrophy, occurs mostly during the early growth stage where protocorms are still achlorophyllous and may continue until adulthood in various genera (Rasmussen et al. 2015; Yeh et al. 2019).

Most orchids associate with *Rhizoctonia-*like fungi, a broad term for several genera resembling the anamorphic morphology of *Rhizoctonia*. As symbionts, there are three genera in this group: *Ceratobasidium, Serendipita* (often associated with *Sebacina sensu lato*), and *Tulasnella* (Rasmussen et al. 2015; Yeh et al. 2019; Pujasatria et al. 2020). Originally, those fungi are saprophytic, endophytic, or even pathogenic. However, among those genera, many species were isolated from orchid roots and proved to be mycorrhizal upon co-inoculation with seeds of respective orchid species (Pujasatria et al. 2020).

Studies on OMF of epiphytic orchids are lacking despite their vast diversity, compared to terrestrial orchids (Yukawa et al. 2009). While the roots of terrestrial orchids are usually subterranean, epiphytic orchids are commonly found growing in exposed environments and are prone to desiccation due to weather change (Rachanarin et al. 2018; Freitas et al. 2020). This leads to a presumably different fungal community compared to those of the underground. Microclimate change and phorophyte architecture also affect the establishment of epiphytic orchids (Rasmussen and Rasmussen 2018), especially for those living in a fluctuating climate, such as temperate regions. The availability of OMF on a particular substrate–in this case, arboreal–is also dependent on such a microenvironment since fungus intrinsically prefers unexposed, moist conditions to grow. When orchid seed lands on the substrate, its germination is not guaranteed since colonization of appropriate OMF is required. Thus, the interaction between ‘the orchid and the fungal community would be important during the orchid’s establishment and growth.

Most epiphytic orchids studied tend to associate with either *Ceratobasidium* or *Tulasnella* (Yukawa et al. 2009; Zettler et al. 2013; Rachanarin et al. 2018; Pujasatria et al. 2020; Freitas et al. 2020). However, more specifically, studies on OMFs of monopodial epiphytic orchids are limited. Several studies were already conducted on angraecoids (Hoang et al. 2017; Kendon et al. 2020), either leafy or aphyllous, but still limited to vandaceous (Carlsward et al. 2006).

This study focused on *Vanda* (syn. *Neofinetia*) *falcata*. This epiphytic orchid is known as one of Japan’s ornamental orchids. Naturally, this orchid is distributed across central to southern Japan and reportedly grows on evergreen or deciduous trees (Suetsugu et al. 2015).

Unlike typical epiphytic orchids concentrated in tropical regions, *V. falcata* is adapted to subtropic and to the temperate areas, which may affect its association with fungi on phorophytes. Previous studies suggested that vandaceous orchids–especially those of subtribe Aeridinae–mainly associated with *Ceratobasidium*, proven by metabarcoding and seed germination results (Otero et al. 2002; Yukawa et al. 2009; Hoang et al. 2017; Mújica et al. 2018; Rammitsu et al. 2019; Kendon et al. 2020). Although *Ceratobasidium* strains have also been isolated from *V. falcata* (Rammitsu et al. 2021), mycobionts for seed germination have never been reported. Thus, this study analyzed the whole fungal community in seeds and root samples of *V. falcata* using amplicon sequencing. Additionally, pure fungal cultures were isolated from the same root pieces. Based on microscopic and molecular analysis, it was found that some of the isolated fungi were *Rhizoctonia-*like fungi. Finally, co-inoculation experiment was conducted to determine the interaction of these fungi for their colonization with seeds and roots of *V. falcata*.

## Materials and Methods

### Plant materials

*Vanda falcata* plants growing on a persimmon (*Diospyros kaki*) trunk were obtained from Kihoku Town, Mie Prefecture, Japan (34°06’27“N, 136°14’17” E). The residents eventually cut the trunk due to safety reasons, and it was maintained in Suzuka City of the same prefecture (34°51’05“ N, 136°36’123” E) (Fig. 1A). Roots of young (small to medium plants that never flowered) and adult (those already flowered at least once) plants (Fig. 1B, C) attached to bark were randomly collected, and the harvested samples were stored at 4°C until use. The baiting method assessed fungal diversity inside the seeds: 3 x 6 cm nylon mesh packs containing ca. 100 *V. falcata* seeds were tied adjacent to twelve randomly selected plants, including young and adult plants. The baiting method was conducted during spring (March–June 2020). The seeds and young/adult plant roots were surface sterilized and stored in 70% ethanol at 4°C until DNA extraction.

**Fig. 1.**
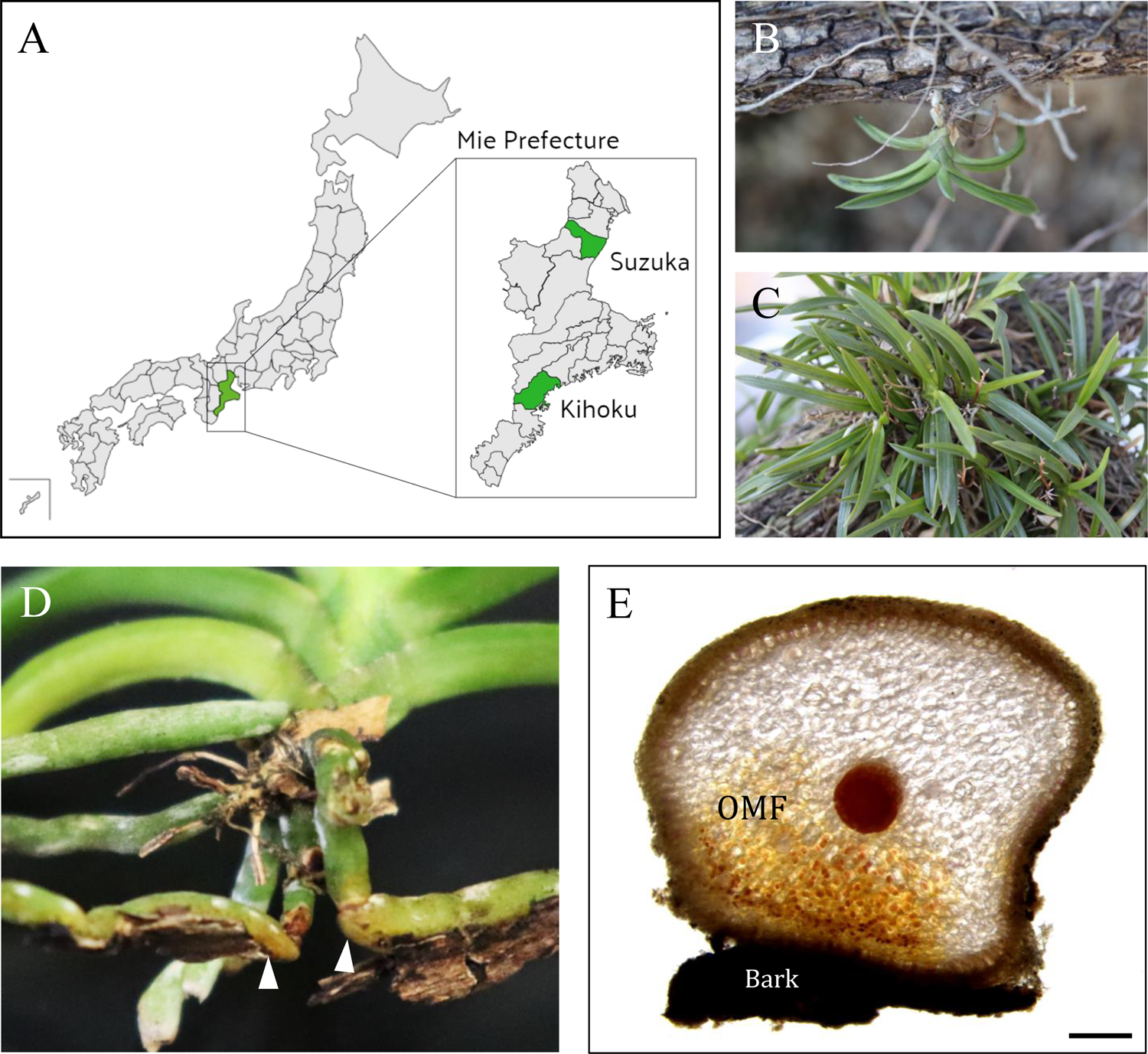
Plant samples. A) Location of Mie Prefecture, Kihoku Town, and Suzuka City (green) inside Japan. B) *Vanda falcata* young plant example. C) Adult plants with old, drying inflorescences. D) Colonized root parts conspicuously turned yellow upon orchid mycorrhizal fungi (OMF) colonization (arrowheads). E) OMF mostly colonizes the root part that attaches to the bark. Bar = 1-mm.

### DNA extraction from plant materials and amplicon sequencing

Seeds isolated from the baiting method and root segments of randomly selected young and adult plants were used as materials for DNA extraction using the Real Genomics Plant DNA Extraction kit (RBC Bioscience, Taipei, Taiwan). After extraction, DNA concentration was measured using DS-11 spectrophotometer (DeNovix, DW, USA). The internal transcribed spacer (ITS) region was amplified using ITS1F_KYO1/ITS2_KYO2 for ITS1 and a pair of gITS/ITS4 for ITS2 (Ihrmark et al. 2012; Toju et al. 2012) (Table S1). Each reaction mixture (20-µl) contains 1-ng genomic DNA, 1-µl of 20-µM primer forward/reverse, and 10-µl KOD One PCR Master Mix (Toyobo, Osaka, Japan). For amplification, PCR was coonducted on T100 thermal cycler (Bio-Rad, CA, USA) using the following program: initial denaturation at 94°C for two minutes, 94°C for 30 secs, followed by 25 cycles of 50°C for 30 secs, 72°C for one minute, and final elongation step at 72°C for seven minutes. PCR results were confirmed using 1% (w/v) agarose gel electrophoresis. The sequencing of libraries prepared by PCR with barcode-containing primers using Illumina MiSeq platform (2 × 300 bp) and data analysis was conducted by the Bioengineering Lab Co., Ltd. (Atsugi, Japan). The raw sequence data obtained as fastq file was processed using the FASTX toolkit v0.0.13 (Hannon 2010) and Sickle v1.33 (Joshi and Fass 2011) to remove adaptor and primer sequences, ambiguous reads, low-quality sequences (quality score less than Q20), and reads no more than 40-bp. Quality-filtered sequences were merged using FLASH v1.12.11 (Magoč and Salzberg 2011) and used for the production of the representative sequences and operational taxonomic unit (OTU) table using Qiime2 (v2020.6) (Bolyen et al. 2019) after removing the illusion and noise sequences using dada2 plugin. The representative sequences were compared to OTUs (97%) in UNITE v8.2 (Nilsson et al. 2019) to classify by taxon using the fitted classifier of Qiime2. Additionally, alpha and beta diversity analyses were conducted using Qiime2 diversity plugin.

Amplicon sequencing results were visualized using RStudio v4.0.2. The *ggpubr* package was used to visualize fungal diversity indices (OTU abundance, Chao1, Pielou, and Shannon Evenness Index/SEI), and the proportion of each fungal phylum observed in samples (Kassambara 2018). Principal coordinate analysis (PCoA) plots–based on Bray-Curtis index, Euclidean distance, and without any data transformations–were visualized using the *vegan* package (Oksanen et al. 2007). Additionally, analysis of similarity (ANOSIM) was used to calculate PCoA statistical significance (p < 0.05), using the same package.

### Fungi-colonized root morphology observation

Crossing manual sections of fungi-colonized root were stained using 3% (w/v) acid fuchsin in glacial acetic acid (Gange et al. 1999) or UV autofluorescence after clearing in 10% KOH for one hour at 90°C to observe the presence of OMF and other fungi. The samples were then observed under a light microscope (BX53; Olympus, Tokyo, Japan), and images were taken using the equipped digital camera (DP27; Olympus). Statistics for quantification of OMF colonization in roots was calculated using ANOVA based on the results of the Kolmogorov-Smirnov normality test and Bartlett test using *ggpubr* package of RStudio v4.0.2 (Kassambara 2018). Subsequent Tukey *post hoc* test was conducted using the same software.

### Rhizoctonia-like and pathogenic fungi isolation

Roots of young and adult plants were sectioned manually and observed using light (BX53) or stereo microscope (SZX16; Olympus). Subsequently, root sections with fungal colonization in cortex were put on either potato dextrose agar (PDA), one-sixth strength Czapek-Dox agar, or Fungal Isolation Medium (Zettler et al. 2013) supplemented with 10-µg/mL tetracycline. Fungi with *Rhizoctonia-*like characteristics (Oberwinkler et al. 2013; Weiß et al. 2016) were cultured on PDA, malt extract agar (MA), full-strength V8 juice agar (V8), or oatmeal agar (OMA) media at 25°C, and then stored at 4°C for long-term storage, while all non-*Rhizoctonia-*like fungi were cultured only on OMA at 25°C. Hyphae were stained using 0.01% ethidium bromide in 25% ethanol (Singh and Kumar 1991), and the cell wall was stained using a mixture of 1-mg/L calcofluor white and 0.5-mg/L Evans blue, to count the nuclei of *Rhizoctonia-*like fungal hyphae. Samples were observed under a fluorescence microscope (DM2500; Leica, Wetzlar, Germany) equipped with a digital camera (DFC310 FX; Leica) and visualized using ImageJ v.1.53a.

Fungal genomic DNA was extracted from the cultured fungi using the sorbitol-cetrimonium bromide (CTAB) combination method (Inglis et al. 2018). After extraction, DNA concentration was measured as described above. The ITS region was amplified using a combination of ITS1OF/ITS4 (Table S1) (Taylor and McCormick 2008; Toju et al. 2012). The PCR mixture for each sample contains 1-µl fungal genomic DNA, 1-µl of 20-µM primer forward/reverse, and 10-µl GoTaq Green Master Mix (Promega, WI, USA). Amplification was conducted using the following program: initial denaturation at 95°C for ten minutes, followed by 35 cycles at 95°C for 20 secs, 50°C for 30 secs, 72°C for 20 secs, and a final elongation step at 72°C for seven minutes. Successfully amplified DNA fragments confirmed by 1% (w/v) agarose gel electrophoresis were cloned using TOPO TA Cloning Kit (ThermoFischer Scientific, MA, USA) and sequenced using Sanger method with both M13 forward and reverse primers by Eurofins Genomics (Tokyo, Japan). DNA sequences were subjected to BLAST (https://blast.ncbi.nlm.nih.gov/Blast.cgi) to identify the closest taxa.

### Phylogenetic analysis

Fungal ITS DNA sequences were aligned using ClustalW v1.6, and subsequent maximum likelihood (ML) phylogenetic tree construction was conducted using RAxML-HPC v8.2.10 implemented in GENETYX-Windows v15 (Genetyx, Tokyo, Japan) with GTRGAMMA model and 1000 bootstrap analysis to infer the phylogenetic position of fungal OTUs and isolated strains of *Rhizoctonia-*like fungi. The best rooted tree was visualized using Iroki online tool (Moore et al. 2020).

### In vitro seed germination assay

Eleven *Rhizoctonia-*like fungal strains containing *Ceratobasidium, Serendipita,* and *Tulasnella* (Table 1), obtained from culture stock centers, ATCC (American Type Culture Collection), RIKEN bioresource center (JCM strain), NARO Genebank (MAFF strains) and NITE biological resource center (NBRC strains), were inoculated on OMA for two weeks before symbiotic germination assay. Dehiscent capsules formed from naturally pollinated flowers were used to obtain the *V. falcata* seeds. Seeds were sterilized using 3% hydrogen peroxide and washed two to three times with sterilized water. Cultures were stored at 25°C for eight weeks in darkness. Germination stage description (Zettler et al. 2007) was modified according to the preliminary germination assay result (Table 2, Fig. S1). Protocorm growth index (GI) was evaluated using the equation *GI = ∑^n^_i=5_i(S_i_) / ∑ S*, with i; germination stage number (starting from stage 0 to 5), n; total seeds in one treatment, and *n_i_*; total seeds reaching germination stage *i* (Guimarães et al. 2013), and symbiotic cells were counted after six weeks. Each fungal treatment was replicated four times, with each plate containing ca. 100–300 seeds. Additionally, two ascomycetous fungi isolated from roots, *Pyrenochaetopsis* and *Fusarium,* were also subjected to similar germination assay to observe their interaction with seeds (Table 3). Protocorms were stained using 1-g/L calcofluor white counterstained with 0.5-g/L Evans blue, and fungal hyphae were stained using 1-mg/L WGA-Alexa Fluor 594 (ThermoFisher Scientific). Visualization was conducted using a fluorescent microscope (DM2500) equipped with a digital camera (DFC310 FX) and ImageJ v.1.53a, while symbiotic cells were counted using a tally counter from 25 randomly selected protocorms. Based on the results of the Kolmogorov-Smirnov normality test and Bartlett test, the Kruskal–Wallis test was conducted using *ggpubr* package of RStudio v4.0.2 (Kassambara 2018). Subsequently, Dunn’s *post hoc* test with Benjamini-Hochberg false discovery rate adjustment (p < 0.05) was conducted using the *FSA* package of the same software (Ogle et al. 2021).

**Table 1.** *Rhizoctonia*-like fungi used for *Vanda falcata* seed symbiotic germination

| Treatment | Species name | Strain no. | Source |  |
| --- | --- | --- | --- | --- |
|  |  |  | Host plant | Location |
| C.gra53 | *Ceratobasidium gramineum* | MAFF235853 | *Agrostis sp.* * ** | Japan |
| C.sp.87 | *Ceratobasidium sp.* | MAFF244587 | *Dactylorhiza aristata* | Japan |
| C.sp.BI | *Ceratobasidium sp.* | N/A | this study | Japan |
| C.sp.DE | *Ceratobasidium sp.* | N/A | this study | Japan |
| C.sp.TA1 | *Ceratobasidium sp.* | NBRC109234 | *Taeniophyllum glandulosum* | Japan |
| C.sp.TA2 | *Ceratobasidium sp.* | NBRC109235 | *Taeniophyllum glandulosum* | Japan |
| S.ind27 | *Serendipita indica* | ATCC DSM11827 | *Prosopis juliflora* and *Zizyphus nummularia* * *** | India |
| S.ver30 | *Serendipita vermifera* | MAFF305830 | *Cyrtostylis reniformis* | Australia |
| T.asy08 | *Tulasnella asymmetrica* | MAFF305808 | *Thelymitra epipactoides* | Australia |
| T.cal05 | *Tulasnella calospora* | MAFF305805 | unspecified | Australia |
| T.irr96 | *Tulasnella irregularis* | JCM9996 | *Dendrobium dicuphum* | Australia |

\* Non-orchidaceous plants, \*\* isolated as pathogen, \*\*\* isolated as endophyte

**Table 2.** Seed germination stage morphological descriptions

| Stage | Description |
| --- | --- |
| 0 | No response |
| 1 | Imbibition, OMF starts to colonize |
| 2 | Swelling, testa rupture, formation of rhizoids |
| 3 | Formation of shoot primordium |
| 4 | Formation of first crest |
| 5 | Formation of first leaf |

**Table 3.** Isolated fungi from *Vanda falcata* seedling and adult plant roots

| Isolate | Source sample | Closest match in Genbank | Similarity |
| --- | --- | --- | --- |
| BI-103P | Young root | AB507066.1 uncultured Ceratobasidiaceae from *Sedirea japonica* (Miyazaki Prefecture, Japan) | 97.74% |
| DE-52P | Adult plant root | AB507066.1 uncultured Ceratobasidiaceae from *Sedirea japonica* (Miyazaki Prefecture, Japan) | 98.79% |
| MI-3 | Young root | NR_155625.1 *Parateratosphaeria karinae* holotype CBS128774 (South Africa) | 84.20% |
| MI-4 | Young root | NR_153559.1 *Atrocalyx bambusae* holotype MFLU 11-0150 (Thailand) | 97.25% |
| MI-5 | Young root | NR_159064.1 *Subulicystidium harpagum* isotype KAS L 1726a (Réunion Island) | 89.26% |
| NE-4 | Adult plant root | NR_160059.1 *Pyrenochaetopsis microspora* type material CBS 102876 (Montenegro) | 99.81% |
| NE-5 | Adult plant root | NR_137617.1 *Fusicolla violacea* holotype CBS:634.76 (Iran) | 80.92% |

## Results

### Amplicon sequencing

Genomic DNA was obtained from three kinds of samples: seeds, roots of young plants, and roots of adult plants. The seeds were taken from twelve seed packs and combined as one even though no germinating seeds were found. For root samples, three to four root sections (ca. 5-cm each) were cut from a young or adult plant. In root samples, the most conspicuous feature of the symbiotic region was a yellowish root surface that contains digested pelotons (Fig. 1D). This was primarily obvious when the roots were wet. This coloration is unique to the root parts directly attached to *D. kaki* bark (Fig. 1E).

The ITS1 and ITS2 PCR products were subjected to amplicon sequencing by Illumina MiSeq. Each sample sequencing generated 36,027 to 72,301 raw reads for ITS1 and 40,033 to 72,014 raw reads for ITS2. 83% and 89.47% of the whole reads were processed and used for further analysis (Table S2). Phylogenetic OTUs were generated at 97% similarity cutoff using Qiime2 and ranged from 155 to 411 OTUs for each sample in ITS1 and 174 to 476 OTUs for each sample in ITS2 (Table S3, S4). In total, 1274 and 1358 phylogenetic OTUs were obtained for ITS1 and ITS2, respectively.

### General fungal diversity and OMF localization in roots

PCoA results conducted using generated OTUs from both ITS sequenced data showed distinct groupings of fungal communities among the three development stages–seed, young plant, and adult plant (Fig. 2), but ANOSIM did not give a significant result for ITS2 (p < 0.05). R-value of 0.3913 (sig. = 0.0419) for ITS1 and 0.2047 (sig. = 0.1965) for ITS2 shows that the fungal compositions between the stages were not distinguished (Clarke 1993) (Fig. 2). R-value between ITS1 and ITS2 communities has different significance level but both values imply that the degree of discrimination is not changed. The community structure similarities were also supported by Pielou evenness (J’) and SEI, where fungal diversity in all samples was not significantly different (Fig. S2). Ascomycota and Basidiomycota were the most prevalent phyla at the phylum level, with coverage of above 50% in each sample (Fig. S3). The other phyla were Chytridiomycota, Mortierellomycota, and Mucoromycota, which were found in smaller amounts.

**Fig. 2.**
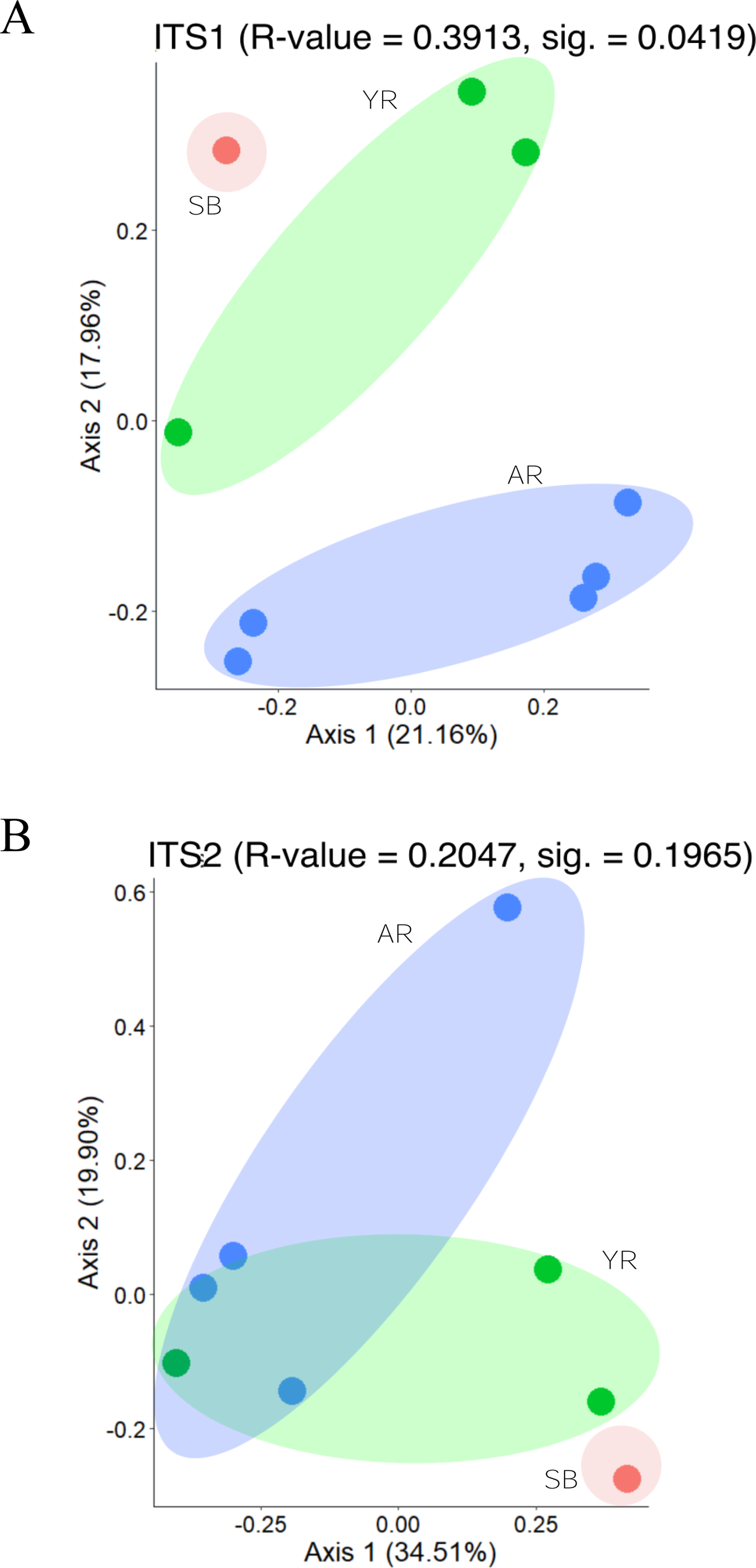
Beta diversity of the fungi in *Vanda falcata* based on the ITS1 and ITS2 sequences. Principal coordinate analysis (PCoA) plots of ITS1 (A) and ITS2 (B) based on Bray-Curtis distance showing a composition of fungal community structure in all samples, e.g., seeds from seed baiting (SB, red), young plant roots (YR, green), and adult plant roots (AR, blue). R-value was calculated using ANOSIM (p < 0.05).

In *Rhizoctonia*-like fungi, *Ceratobasidium* was found in trace amounts in seed samples but was unexpectedly found only in one young root sample. It was also found in two other adult root samples but only in trace amounts. Another similar case is in *Serendipita*, which was relatively abundant only in seed samples. *Tulasnella* was very limited in seed samples but gradually increased in young and adult root samples (Fig. 3A, S4). Thus, while the whole fungal community did not alter, existence of *Rhizoctonia-*like fungi (*Ceratobasidium, Serendipita,* and *Tulasnella*) tended to change over time following the growth and development of *V. falcata*.

**Fig. 3.**
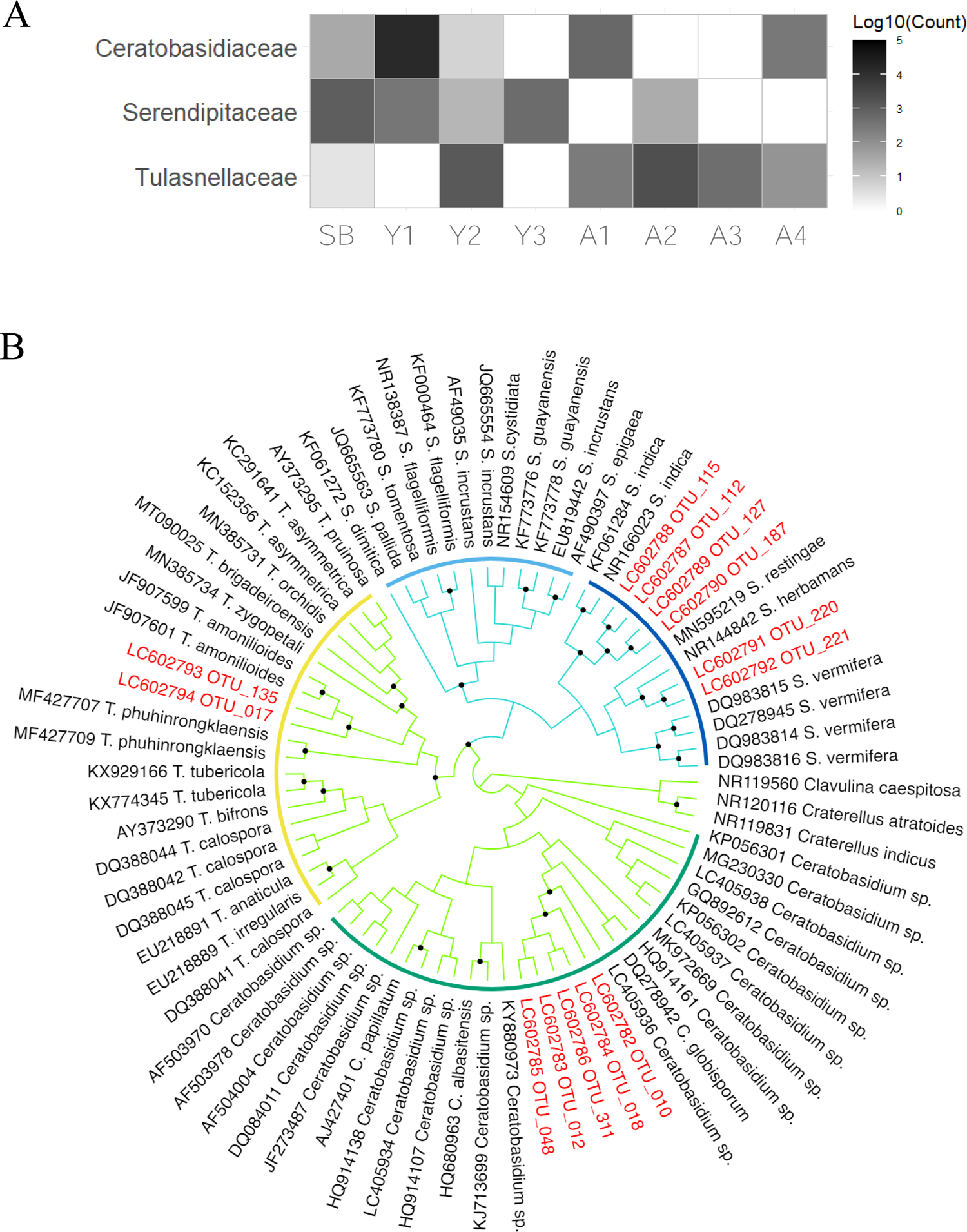
*Rhizoctonia*-like fungal OTUs inferred from amplicon sequencing results. A) Heat map diagram showing *Rhizoctonia*-like fungi OTUs present in seed (SB), seedling (Y1-3), and adult (A1-4) root samples. B) Maximum likelihood (ML) phylogenetic tree of OTUs assigned to each order (colored branches)–Cantharellales (chartreuse) and Sebacinales (light blue)–and family (arcs): Ceratobasidiaceae (green), Serendipitaceae (blue), Sebacinaceae (light blue), and Tulasnellaceae (yellow). Only bootstrap numbers above 80% (dotted nodes) are shown. *Clavulina caespitosa, Craterellus atratoides,* and *Craterellus indicus* are used as outgroups from Cantharellales.

Both ITS sequences exhibited relatively similar diversity indices. However, ITS2 has been considered a suitable marker for revealing the operational taxonomic richness and taxonomy specifics of fungal communities due to the broader taxonomic information (Yang et al. 2018). Therefore, this study mainly focused on the ITS2 sequences in subsequent analysis.

### Identification of Rhizoctonia-like fungal OTUs

Among all basidiomycetous fungal sequences, fourteen OTUs annotated as Rhizoctonia-like genera (Table S5) were extracted for the heatmap diagram (Fig. 3A). ML phylogenetic trees based on the representative sequences of each OTU were constructed to see whether the fungi belong to a particular OMF species in their respective families (Fig. 3B). Five *Ceratobasidium* OTUs (accession numbers LC602782–LC602785) were closely related to a mycobiont of epiphytic orchid *Taeniophyllum glandulosum* isolated in Japan (LC405936) (Rammitsu et al. 2019). Six *Serendipita* OTUs (LC602788–LC602792) assigned to Sebacinaceae were divided into two clades inside Serendipitaceae: two OTUs were related to *Serendipita herbamans* (NR144842), an endophyte of *Bistorta vivipara* (Polygonaceae) (Riess et al. 2014) in one clade, and the other four OTUs were close to *S. indica* (NR166023, KF061284, MH863568), a well-known endophyte occurring in several flowering plants (Weiß et al. 2016). Two *Tulasnella* OTUs (LC602793 and LC602794) were related to *T. irregularis* (EU218889) and *T. amonilioides* (JF907601 and JF907599), mycobionts first reported in *Dendrobium affine* (syn. *dicuphum*) and *Encyclia dichroma*, respectively (Warcup and Talbot 1980; Almeida et al. 2014).

### Isolation of Rhizoctonia-like fungi from plant samples

We initially attempted to isolate pure fungal cultures from all samples (seeds, protocorms, and roots) to verify that *V. falcata* associated with *Rhizoctonia*-like fungus during the development *in situ*. However, since no germinating seeds were found, roots of randomly selected young and adult plants were only used for the fungal isolation. We could only isolate two pure isolates of *Rhizoctonia*-like fungi from these root sections despite more than 30 attempts, including those with conspicuous symbiotic regions. We faced two major challenges to obtain a pure culture, which include: (1) most pelotons were already digested, and (2) other endophytic ascomycetous fungi–mostly those morphologically resembling *Trichoderma*, *Cladosporium*, or *Phoma–*often outgrew preferred *Rhizoctonia-*like fungi in the isolation media. In root samples, ascomycetous fungi were found in the epidermis with its conidial form (Fig. 4A). Based on the conidia structure, some of these fungi were identified as *Lasiodiplodia* and Pleosporales (Zhang et al. 2012). However, upon further examination into exodermis, no fungi colonized the exodermal cells. Instead, *Rhizoctonia-*like fungi formed peloton in exodermal passage cells. Based on hyphal morphology, it resembles *Rhizoctonia-* like fungus isolated from roots: both possess hyphae with irregular width, commonly called monilioid hyphae (Fig. 4B, S5). Upon forming peloton inside passage cells, the *Rhizoctonia-* like fungi infiltrate cortical cells to form the peloton network. Additionally, passage cells are concentrated in lower part of the root. Following this, OMF is also mostly found in the same part (Fig. 4C).

**Fig. 4.**
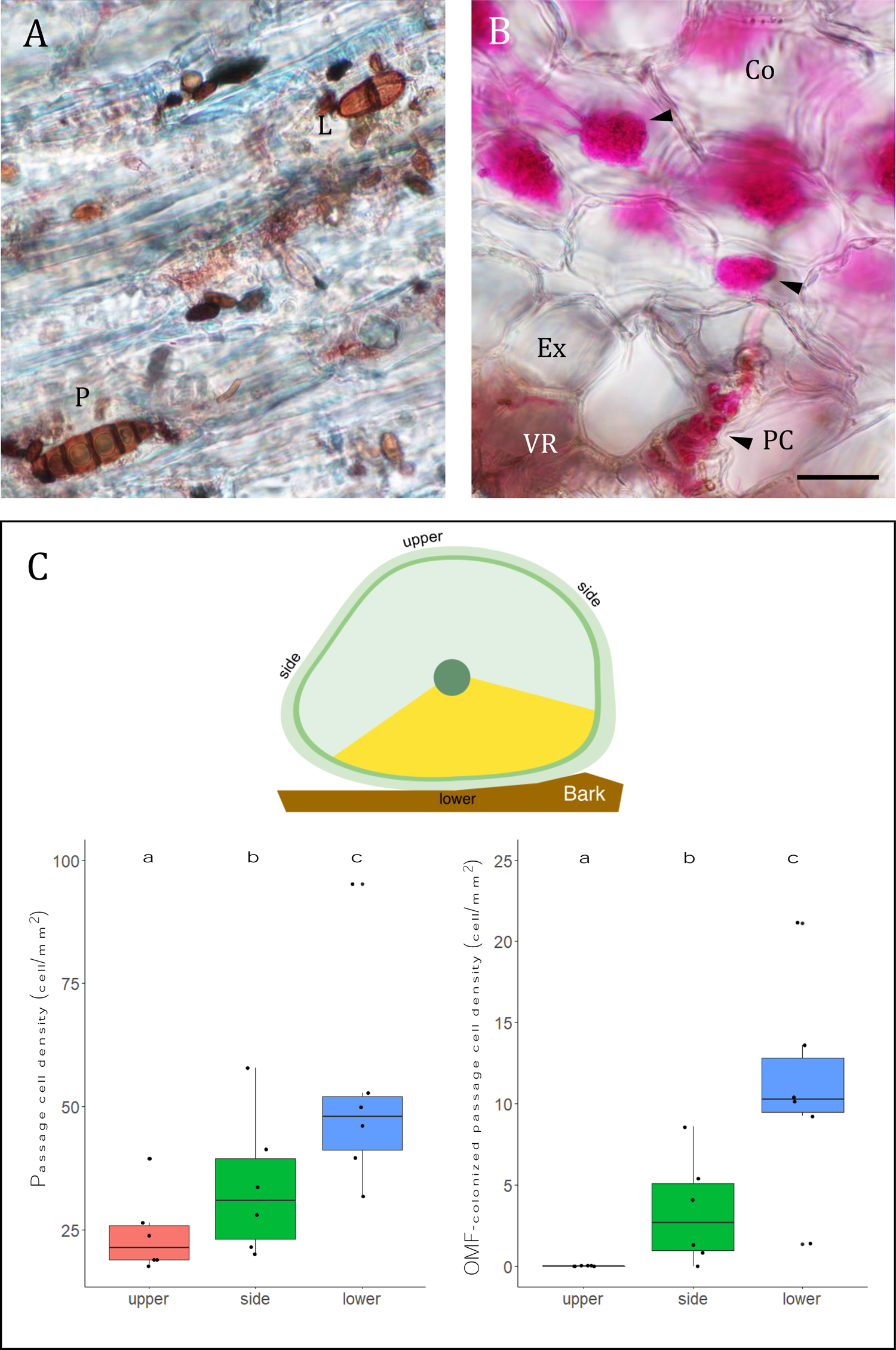
Colonization of ascomycetous and orchid mycorrhizal fungi (OMF) inside roots. A) Conidia of *Lasiodiplodia* (L) and unidentified Pleosporales (P) on the epidermis. B) OMF hyphae (arrowheads) infiltrating the cortical cells (Co) through velamen (VR) and passage cell (PC) of exodermis (Ex). Bar = 40 μm. C) Localization of exodermal passage cells in the upper, side, and lower parts of root (left), and those colonized by OMF (right). Different letters represent significant difference by Tukey’s test at p < 0.05.

The isolated *Rhizoctonia-*like fungi were named BI-103P from young and DE-52P from the adult plant root segments (Table 3, Fig. S6A, B). Morphologically, those fungal strains followed common descriptions for *Rhizoctonia*: branching at a right angle, constriction of hyphae at the site of branching, occasionally visible dolipore septa, and septation at a short distance after branching (Ogoshi 1975). The ITS sequences of these isolates were subjected to pairwise DNA alignment, a feature of Mycobank (https://www.mycobank.org/) and BLAST, which revealed that both strains belong to Ceratobasidiaceae. These strains were closest to AB507066.1, an uncultured *Ceratobasidium* from another epiphytic orchid *Phalaenopsis* (syn. *Sedirea*) *japonica* from Miyazaki Prefecture in southern Japan (Yukawa et al. 2009). We also found that the hyphae are binucleate, indicating that these fungi are traditionally classified into ‘binucleate *Rhizoctonia’* (Fig. S6C, D).

### In vitro seed germination using isolated Ceratobasidium strains and Ascomycota fungi

*V. falcata* seeds were used for symbiotic germination using isolated *Ceratobasidium* strains and other *Rhizoctonia-*like fungi strains including *Serendipita* and *Tulasnella* obtained from culture stock centers (Table 1). Seeds inoculated with compatible OMF started to undergo imbibition and usually formed rhizoids upon entering stage 2 (Fig. S1A). Protocorms started to turn greenish at the onset of stage 3, and shoot primordium was developed as an irregular protrusion on the terminal part (Fig. S1B). This protrusion will eventually form a crest (Fig. S1C). Compared to other stages, stage 4 protocorms were rarely observed. This stage is characterized by the formation of the first leaf (Fig. S1C). Based on fluorescence imaging, fungal colonization started from the suspensor and spread up to 75% of protocorm size (Fig. S1D).

Based on the germination stage description summarized in Table 2, four of six Ceratobasidium strains (C. sp. BI (BI-103P), C. sp. DE (DE-52P), C. sp. TA1, and C. sp. TA2) yielded remarkably higher GI than others based on Kruskal-Wallis test (p < 0.05) (Fig. 5A). Following GI, symbiotic cell count also increased. Protocorms yielded from suitable fungal treatments also had a high symbiotic cell count (Fig. 5B). It was found that these four strains are included in the same phylogenetic clade (Fig. 6). However, few symbiotic cells were found in other Ceratobasidium, Serendipita, and Tulasnella treatments. These strains could only induce germination until stage 1 even after eight weeks. The protocorms kept swollen, but no pathogenic effect was observed, indicating that these strains simply did not further associate with protocorms, and the swelling was merely a result of imbibition. These results indicated that V. falcata seeds may have specificity toward a group of Ceratobasidium fungi.

**Fig. 5.**
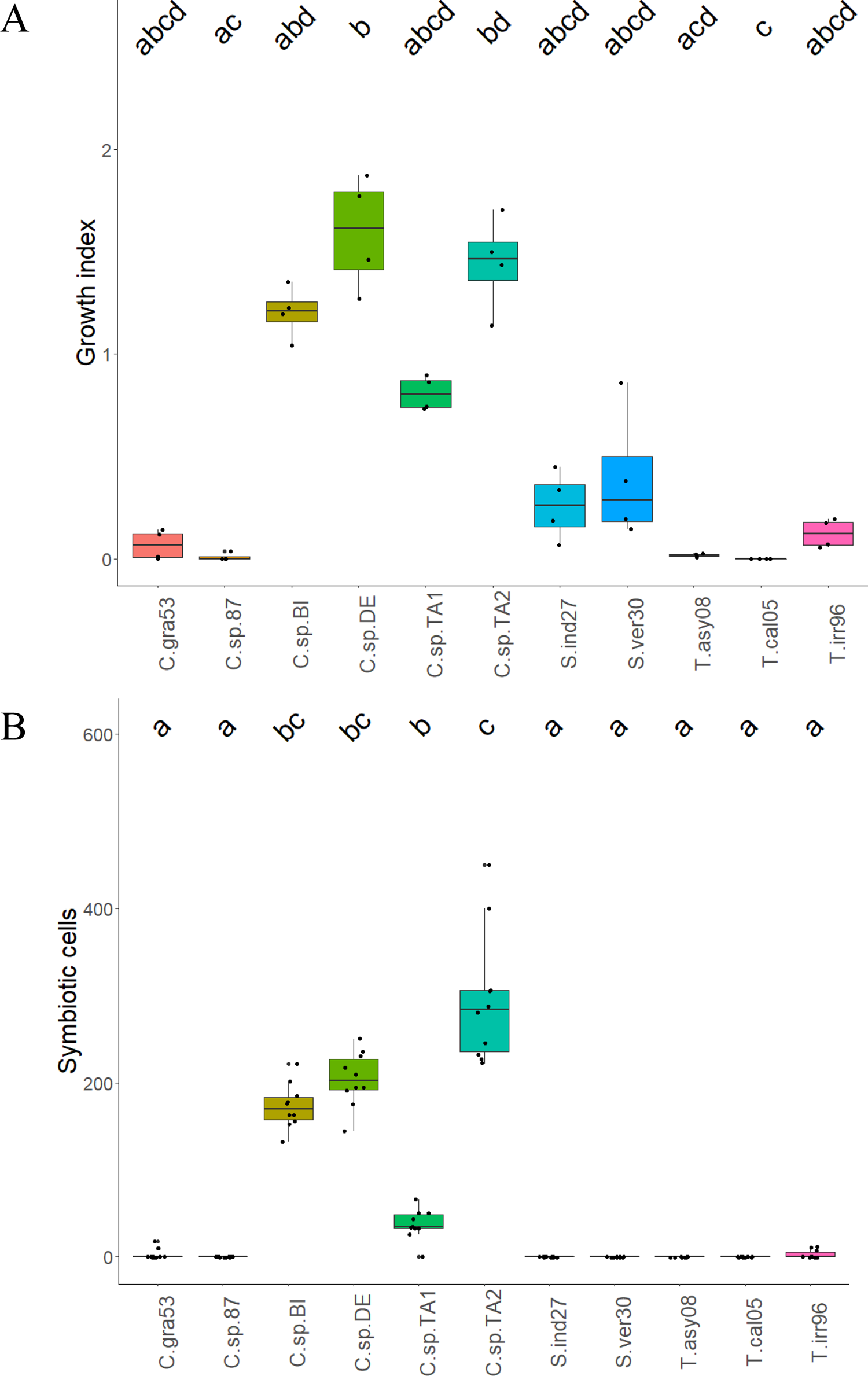
*In vitro* germination assay results. *Vanda falcata* seeds were inoculated with eleven *Rhizoctonia-*like fungi, including those isolated in this study (BI-103P and DE-52P). Boxplots showing growth index (A) and symbiotic cell count (B) after eight weeks. Different letters represent significant difference by Dunn’s test at p < 0.05.

**Fig. 6.**
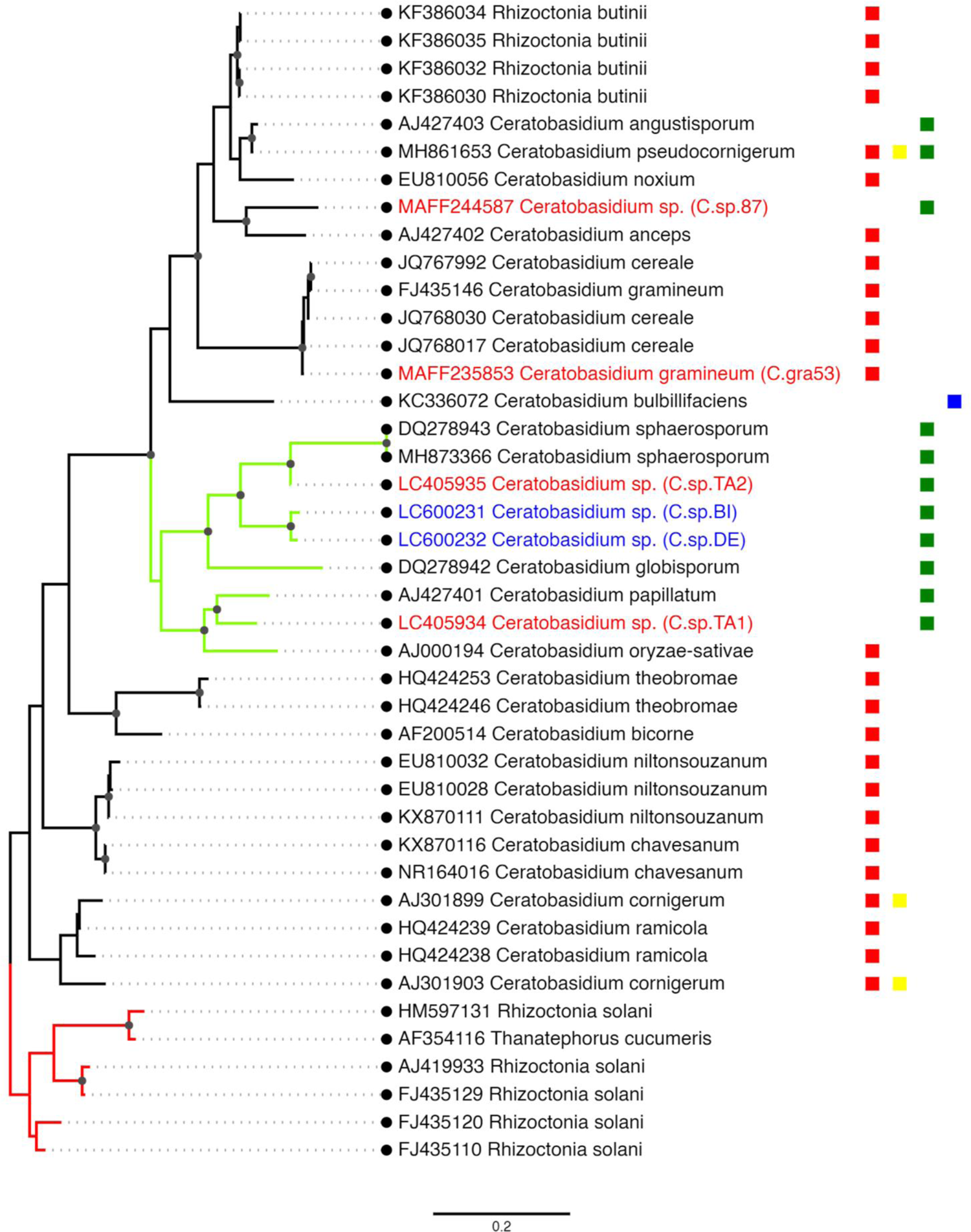
Maximum likelihood phylogenetic tree of *Ceratobasidium* strains, including those inducing seed germination of *Vanda falcata* (red) and other isolated strains (blue). Dotted nodes indicate a bootstrap number of 80% or higher. The clade containing strains with the best germination output is shown in chartreuse branches. Adjacent boxes indicate the nutritional mode of each species, i.e., plant-pathogen (red), saprobe (yellow), orchid mycorrhizal fungus (green), and lichen-forming (blue). *Rhizoctonia solani* strains are used as an outgroup (red branches).

During the process of isolating the *Rhizoctonia*-like fungi from root samples, two Ascomycota fungi *Pyrenochaetopsis* (NE-4) and *Fusarium* (MI-5) (Table 3), were also isolated and identified. Since these are known as pathogenic fungi, the effects of these non-*Rhizoctonia-*like fungi on *V. falcata* seeds were also analyzed using *in vitro* germination assay. At the early stage, seeds were swelling and colonized by *Pyrenochaetopsis* but eventually killed after pycnidia formation, indicated by the blackening of seeds (Fig. 7A). Similarly, the seeds sown on *Fusarium* were heavily colonized and were eventually degraded (Fig. 7B). These results suggested the importance of partitioning between ascomycetous and *Rhizoctonia-*like fungi inside the roots. Additionally, these ascomycetous fungi were incompatible with *V. falcata* seeds for germination.

**Fig. 7.**
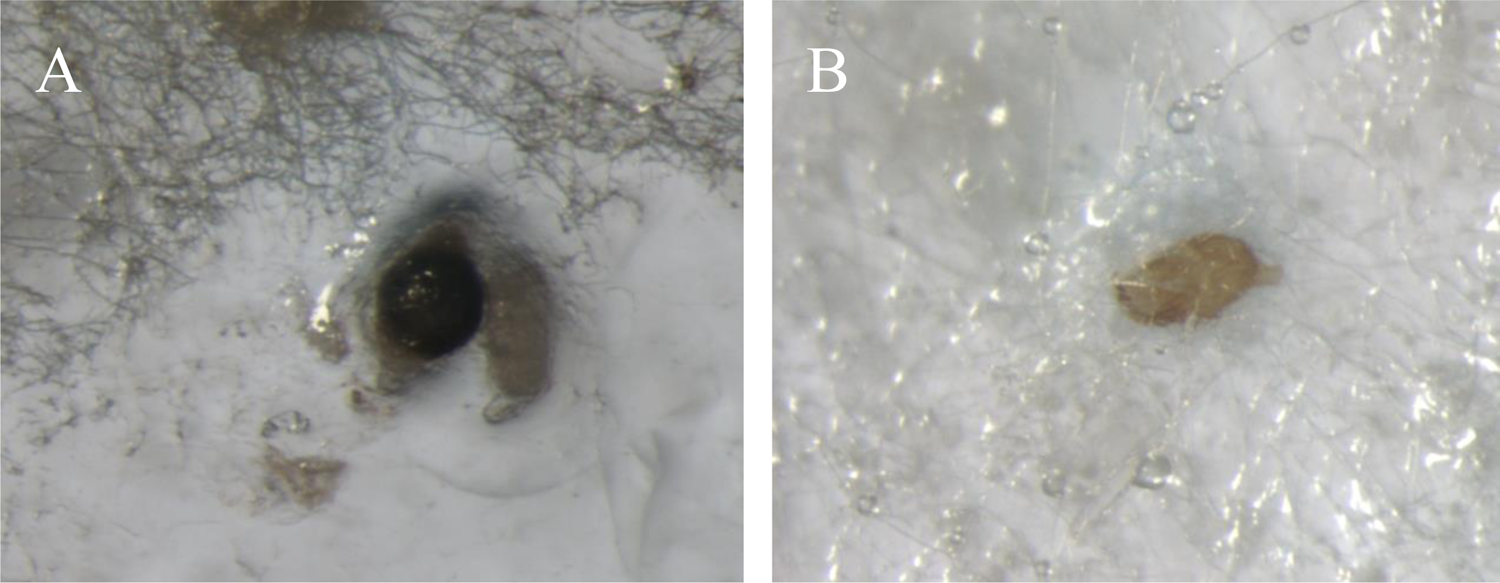
Infected *Vanda falcata* seeds upon inoculation with pathogenic ascomycetous fungi isolated from *V. falcata* roots. A) Seeds infected with *Pyrenochaetopsis.* Black, swollen seed (arrow) was filled with pycnidium. (B) Seed colonized by *Fusarium.* Although inconspicuous, the seed starts to degrade.

## Discussion

While most reports on mycorrhizal associations of the Vandeae tribe come from the Angraecinae subtribe (angraecoids), reports on the Aeridinae subtribe members are still minimal. This study focused on *V. falcata* fungal community structure of the three developmental stages, i.e., seeds, young plants, and adult plants. This study emphasizes this point to show how an orchid maintains its inner fungal community in each developmental stage. All observations provided several findings. Firstly, Ascomycota was the dominant phylum in all examined seeds, young plants, and adult plants. Secondly, ascomycetous, some of which are potential pathogenic fungi, colonized epidermal areas of *V. falcata* roots, while *Rhizoctonia*-like fungi (OMF) colonized the cortical area. Thirdly, *V. falcata* associates with a group of *Ceratobasidium* fungi for seed germination.

Although amplicon sequencing results are not thought to directly reflect fungal community outside samples (i.e., bark surface), the results suggest that the growing substrate does not necessarily provide suitable conditions for OM establishment in epiphytic orchids. Based on the information obtained from PCoA, it was shown that the fungal community in seeds, young, and adult plants were similar with slight differences. Ascomycetous fungal classes, such as Sordariomycetes, Dothideomycetes, and Leotiomycetes frequently occur in all samples, and most of the members are known to be saprobic or parasitic in several habitats, including tree surface where epiphytic orchids coexist (Schoch et al. 2009; Herrera et al. 2010). Seeds landing on bark become the potential hosts of these fungi, especially for *Rhizoctonia*-like fungi, present on the same site. Although it is unknown whether the proportion of ascomycetous and *Rhizoctonia*-like fungi on *D. kaki* bark is similar to that inside *V. falcata* seeds, in the end, ascomycetous fungi will be dominant in the seeds. Even if compatible *Rhizoctonia*-like fungus is present inside the seeds, ascomycetous fungi could encompass the seeds even before germination occurs. Severe infection of *Pyrenochaetopsis* and *Fusarium,* which were isolated from healthy roots upon inoculation with these fungi, indicates that the presence of ascomycetous fungi is a potential hinderance for *V. falcata* germination. As with any endophytic relationship, there is a balanced state between endophyte and roots where the endophyte inhabits within the root without damage to the root. However, when the endophyte was grown on nutrient rich media, it may exhibit virulence against the host (Sarsaiya et al. 2020). In this study, *V. falcata* seeds dispersed on the substrate was vulnerable to *Pyrenochaetopsis* and *Fusarium*.

In the case of *Rhizoctonia-*like fungi, sequence reads in *Serendipita* OTUs were high in seeds, but they much lower in young and adult roots. Conversely, little reads in *Tulasnella* were found in seeds, but its occurrence increases in young and adult roots. The occurrence of *Serendipita* OTUs compared to that of *Ceratobasidium* and *Tulasnella* in seeds indicates the shifting of *Rhizoctonia-*like fungi preference during development but may not necessarily reflect the actual OM association. If *Serendipita* is the OMF of *V. falcata*, seeds in baiting samples should have germinated. Although the number of OTU sequences for *Serendipita*, *Tulasnella*, and *Ceratobasidium* were different at each developmental stage as mentioned, the function of these Rhizoctonia-like fungi is difficult to be speculated based only on the current data and needs to be further investigated.

Among all fungi isolated from root sections, *Ceratobasidium* sp. BI and *C.* sp. DE from young and adult plant roots were confirmed to associate with *V. falcata* due to their capability to propagate seed germination. Interestingly, another strain *Ceratobasidium sp.* TA2 isolated from an epiphytic orchid *Taeniophyllum glandulosum* in Shizuoka Prefecture (Rammitsu et al. 2019) induced better germination results. These three *Ceratobasidium* strains formed a similar clade in the phylogenetic analysis. Therefore, it is suggested that *V. falcata* associates with a narrow range of *Ceratobasidium* in seed germination. Associating with various fungi is advantageous if the OMF occurrence is sporadic, which allows the orchid to readily develop with any compatible fungus available on each growing site (Xing et al. 2019). Generally, it is accepted that orchids with high mycorrhizal specificity are most likely to be rare due to high dependency on OMF distribution in their natural habitats. In the case of *V. falcata*, its distributions were constricted from central to southern Japan with various types of phorophytes (Suetsugu et al. 2015; Rammitsu et al. 2019, 2021). Accordingly, *V. falcata* has high phenotypic variation even for wild plants across Japan, causing such extensive association to be beneficial to the plant for distribution.

We propose that the partitioning of OMF and ascomycetous fungi protects the mycorrhizal root. In this study, ascomycetous fungi mainly detected in *V. falcata* roots were Sordariomycetes, Dothideomycetes, and Leotiomycetes, commonly known as saprobes and pathogens on tree bark (Naranjo-Ortiz and Gabaldón 2019). Primarily based on their saprobic nature, it is plausible that these fungi mainly colonize *D. kaki* bark, and through the orchid root-bark interface, these fungi further penetrate velamen. Although OMF can penetrate into cortical cells, invasion of the ascomycetous fungi is inhibited by exodermis, thus accumulating them inside velamen. Orchid roots typically contain exodermis with lignification of the tangential walls and smaller, non-lignified passage cells that allow OMF to penetrate the cortical cells (Esnault et al. 1994). It is also reported that the passage cell is related to OMF colonization; that is, the root part with denser passage cells has more OMF colonization (Chomicki et al. 2014). Based on these concepts, we also confirmed that passage cells exist in both root surfaces, those attached to bark and those exposed to the air, and the lower part attached to bark had dense passage cells, which was related to higher mycorrhizal colonization in this part. While passage cells are constantly available for OMF to infiltrate, the upper parts of the root mostly remain uncolonized. The exact explanations for this finding are still lacking, but we suggest that chemical (deposition of phenolics, etc.) or environmental (light exposure, etc.) factors might be included. Further studies on how internal and external factors affect epiphytic root colonization by fungi are required to elaborate on this phenomenon.

## Conclusions

To sum up, *V. falcata’s* fungal community structure is similar across growth development. Ascomycota was the dominant phylum, while the others were found in a smaller amount, even for *Rhizoctonia*-like fungi (Fig. 8). It was also confirmed that seeds of *V. falcata* germinated in symbioses with *Ceratobasidium* isolated from its roots and another strain isolated from another orchid. We also propose that the innate regulation of fungal entry also causes this balanced fungal community and partitioning of ascomycetes as well as OMF. Therefore, further studies on how root balances fungal colonization are required to decipher this partitioning mechanism in *V. falcata* or any other epiphytic orchids.

**Fig. 8.**
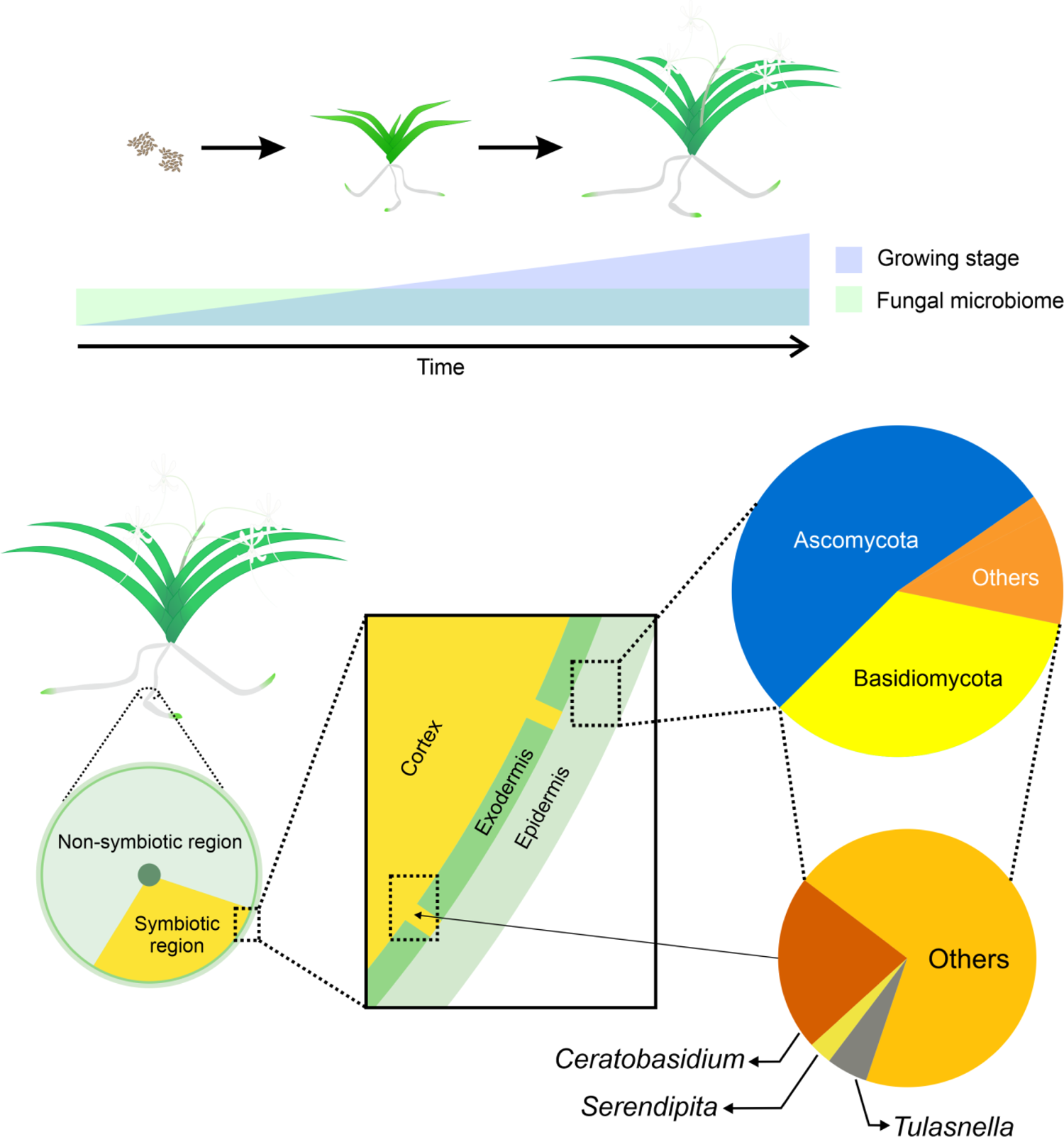
Inner fungal community composition in *Vanda falcata.* Along with growth and development (i.e., from seed germination to reproductive stage), despite an increase in size and trophic mode, the inner fungal community has a similar structure with Ascomycota as the dominant phylum, followed by Basidiomycota. Even for Basidiomycota, *Rhizoctonia-*like fungi abundance is less than half compared to other genera. Among all *Rhizoctonia-*like fungi, *Ceratobasidium* is the OMF of *V. falcata* based on germination assay results, thus correlating with its higher portion. Additionally, only *Rhizoctonia-*like fungus colonizes cortical cells through exodermal passage cells.

## Data Accessibility

The raw sequence reads have been deposited into the DNA Data Bank of Japan (DDBJ) Sequence Read Archive database under the accession number DRA012420 for ITS1 and DRA012422 for ITS2. OTU sequences were registered as LC602782-LC602786 for *Ceratobasidium,* LC602787 - LC602792 for *Serendipita*, and LC602793 - LC602794 for *Tulasnella.* ITS sequences of isolated *Ceratobasidium* were registered as LC600231for BI-103P and LC600232 for DE-52P.

## Author Contributions

GCP, CM, MY, and HK designed the experiments; GCP and IN performed the experiments and analyzed the sequencing data; GCP, CM, MY, and HK wrote the manuscript. All authors approved the final manuscript.

## Acknowledgments

We are grateful to the Japanese Ministry of Education, Culture, Sports, Science, and Technology (MEXT) scholarship to GCP and Research Fellowships of Japan Society for the Promotion of Science (JSPS) for Young Scientists (grant number 201801755) to CM. We also thank Takahiro Yagame (Mizuho Kyodo Museum) for the provision of *Ceratobasidium* TA1-1 and TA2-1, Kenji Suetsugu (Kobe University) for the critical reading of the manuscript, and Rudy Hermawan (Tottori University) for technical assistance.

## Conflict of interest

The authors declare no competing interests.

## Notes

### Competing Interest Statement

The authors have declared no competing interest.

