## Supplemental Figure 1-6 for "Regulation of balanced root fungal community of *Vanda falcata* (Orchidaceae) by partitioning its mycorrhizal fungi and ascomycetous fungi across growth and development"

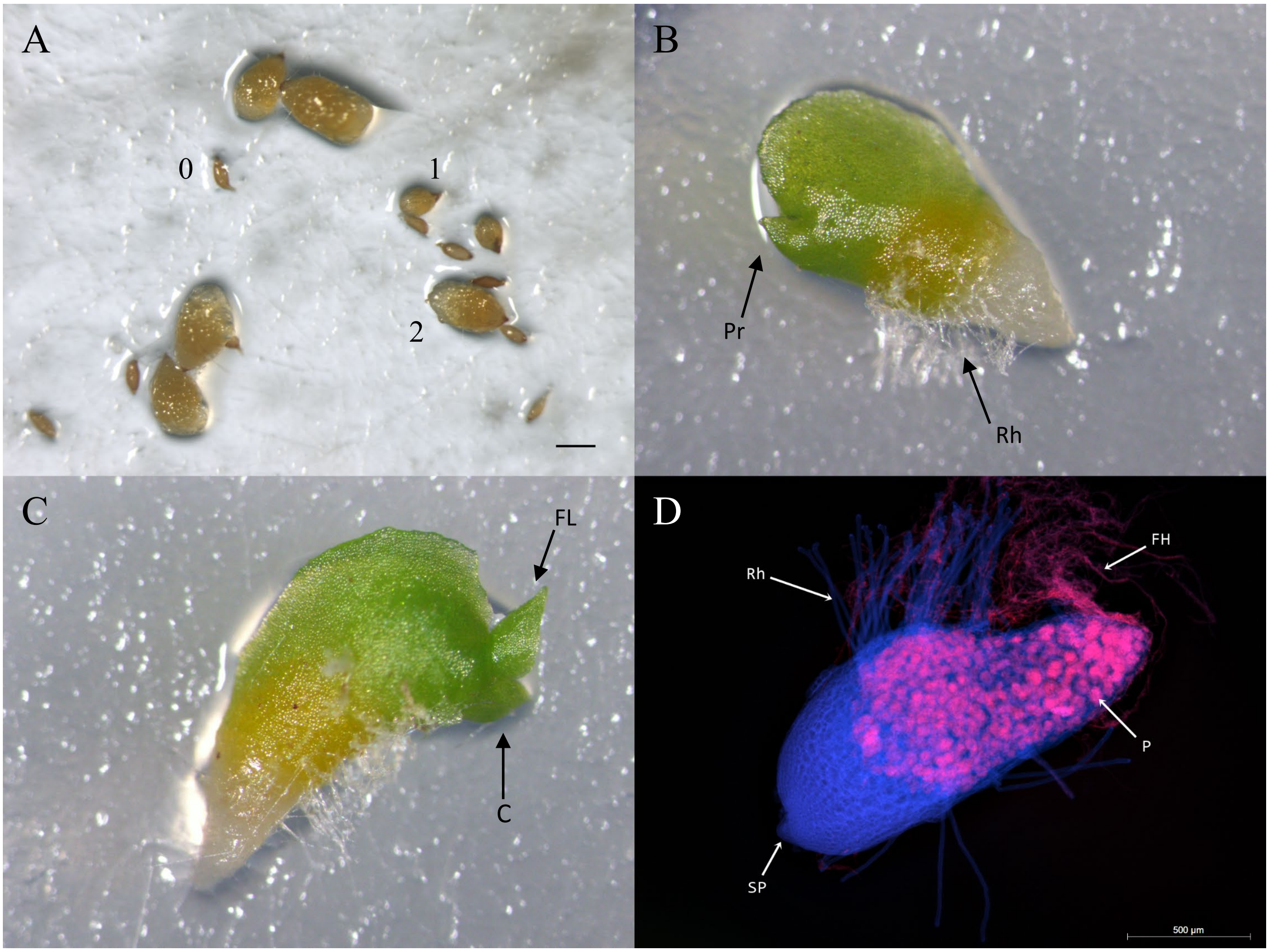


**Fig. S1.** Protocorms at different stages. A) Stage 0, 1, and 2. B) Stage 3 with protrusion (Pr) and rhizoids (Rh). C) Stage 4 with crest (C) and first leaf (FL). D) Fluorescing stage 3 protocorm with peloton (P), free hyphae (FH), rhizoid (Rh), and shoot primordium (SP).

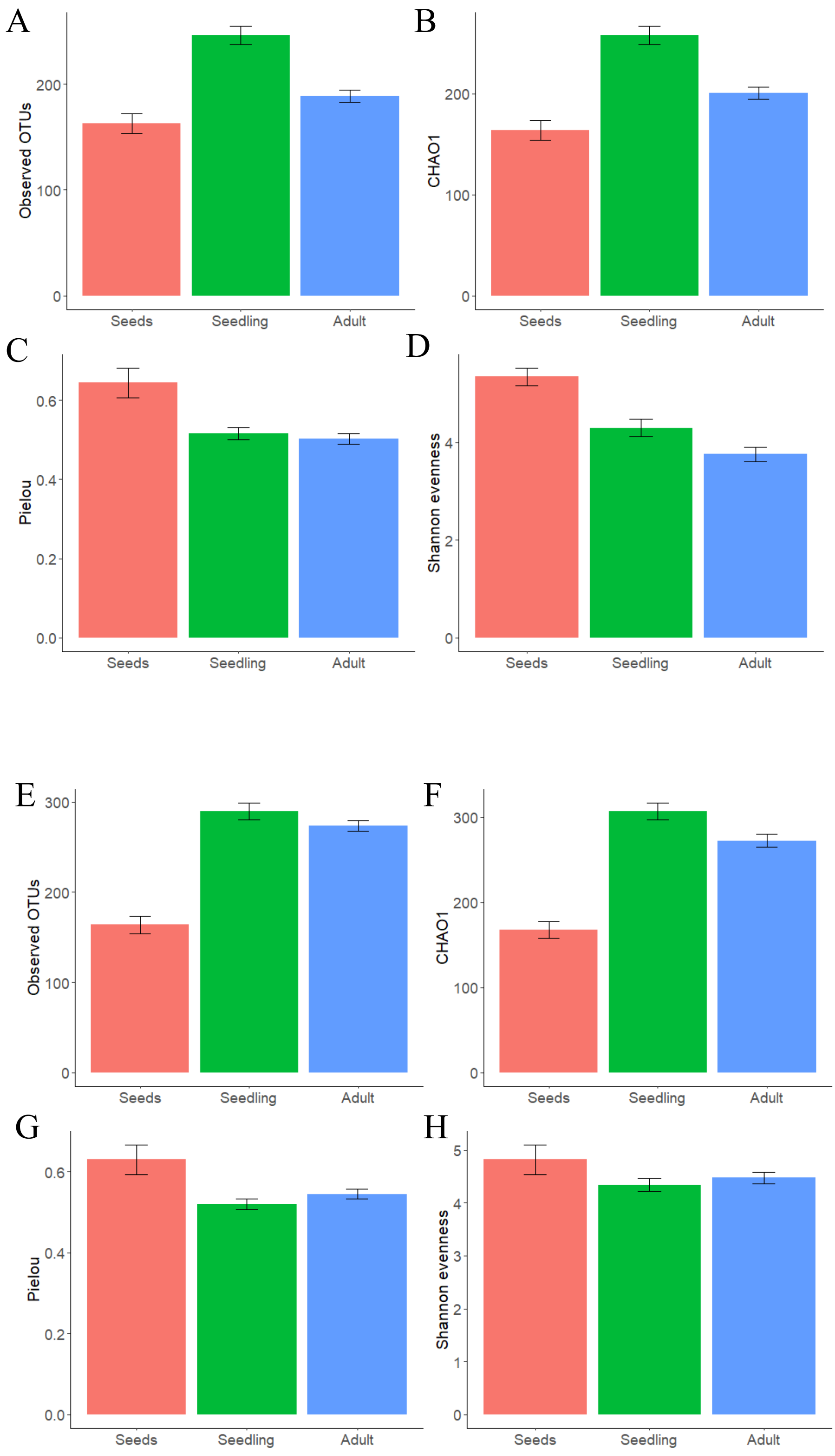


**Fig. S2.** Bar plot of diversity indices (OTU abundance, Chao1, Pielou, and Shannon Evenness Index) of ITS1 (A-D) and ITS2 (E-H).

A

B

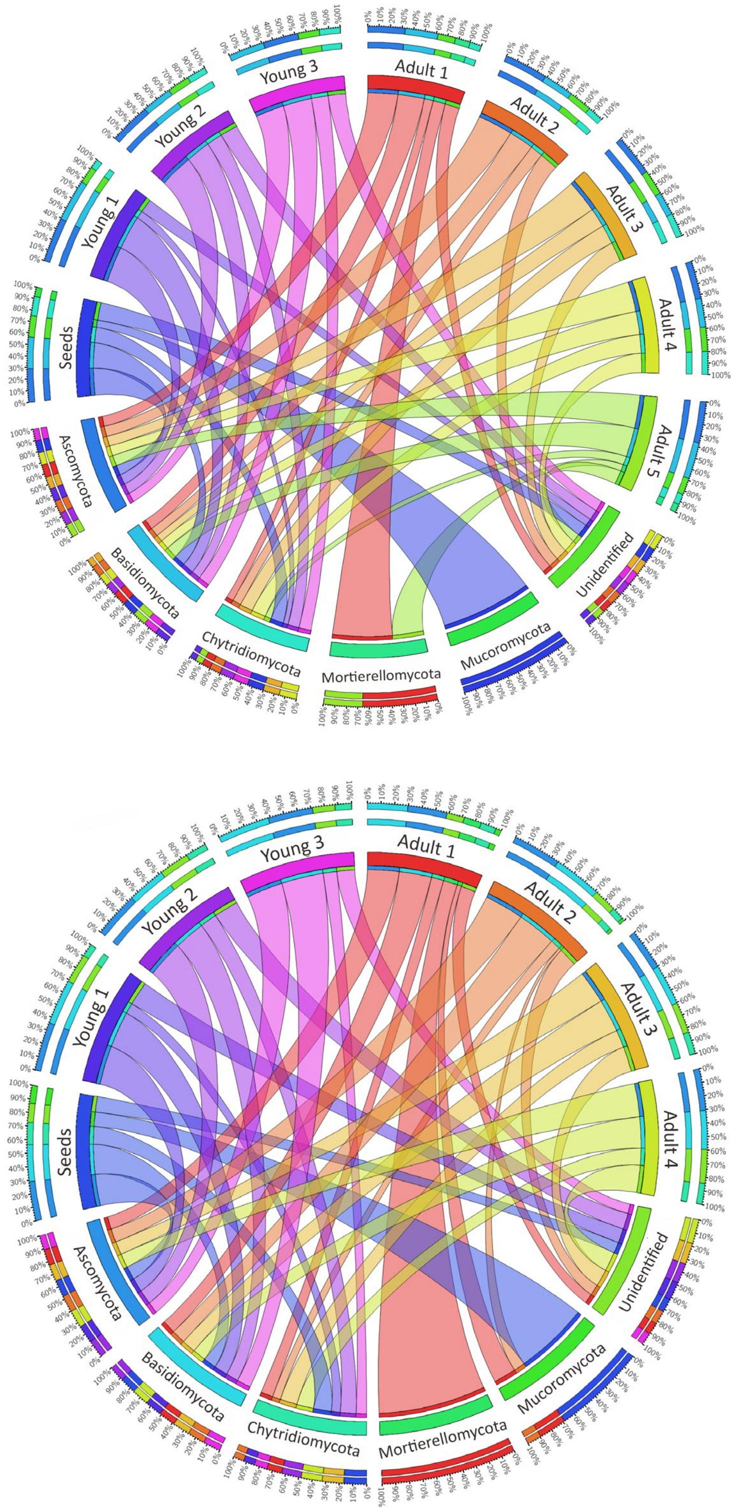


**Fig. S3.** Circos ribbon diagram showing ITS1 (A) and ITS2 (B) fungal phyla present in seed and root samples. Outer circles with percentage shows relative proportion of a family in each sample. Inner circle shows either sample or phylum names.

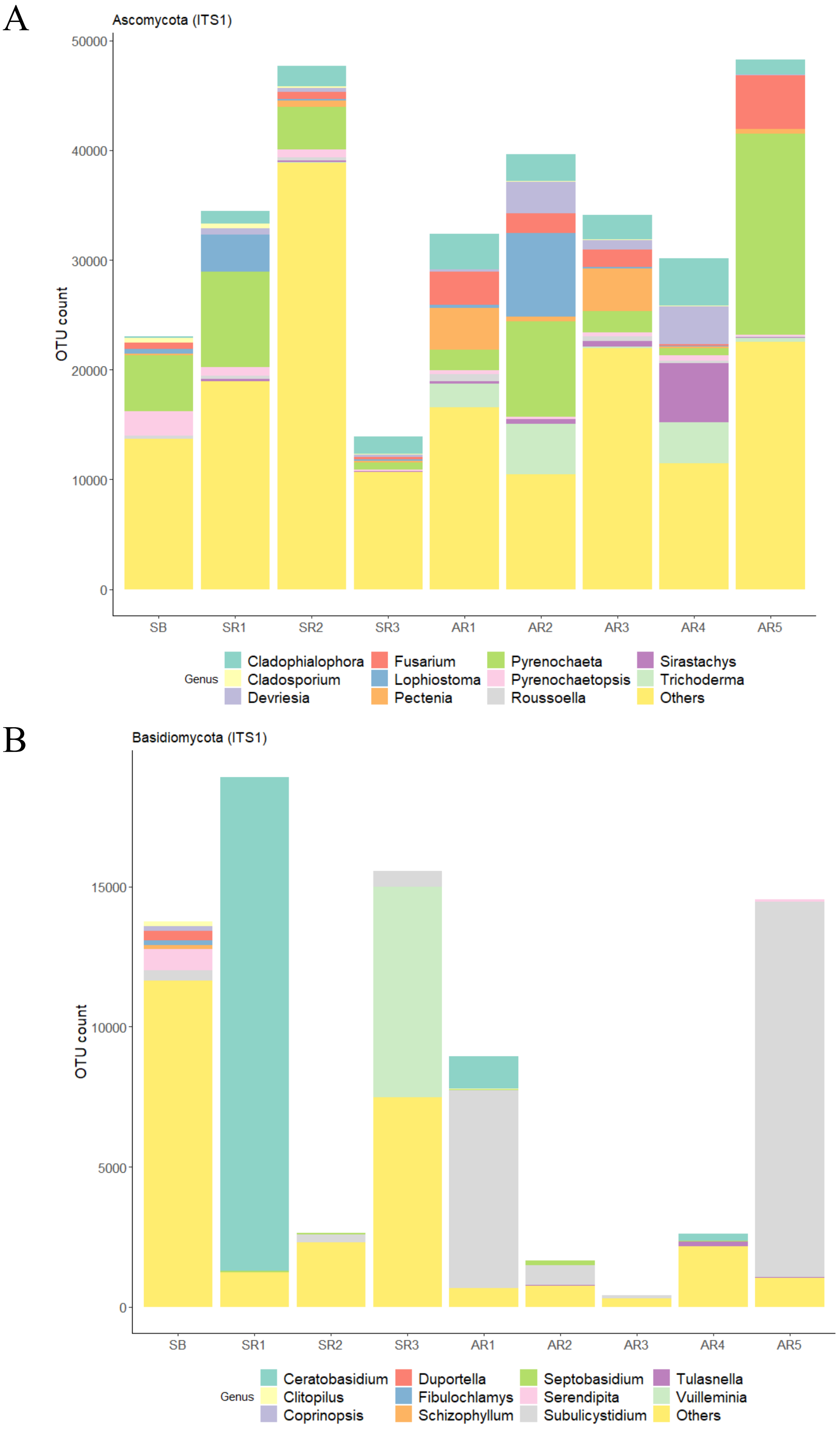


**Fig. S4.** Major Ascomycota and Basidiomycota taxa count in each sample derived from ITS1 (A, B) and ITS2 (C, D).

C

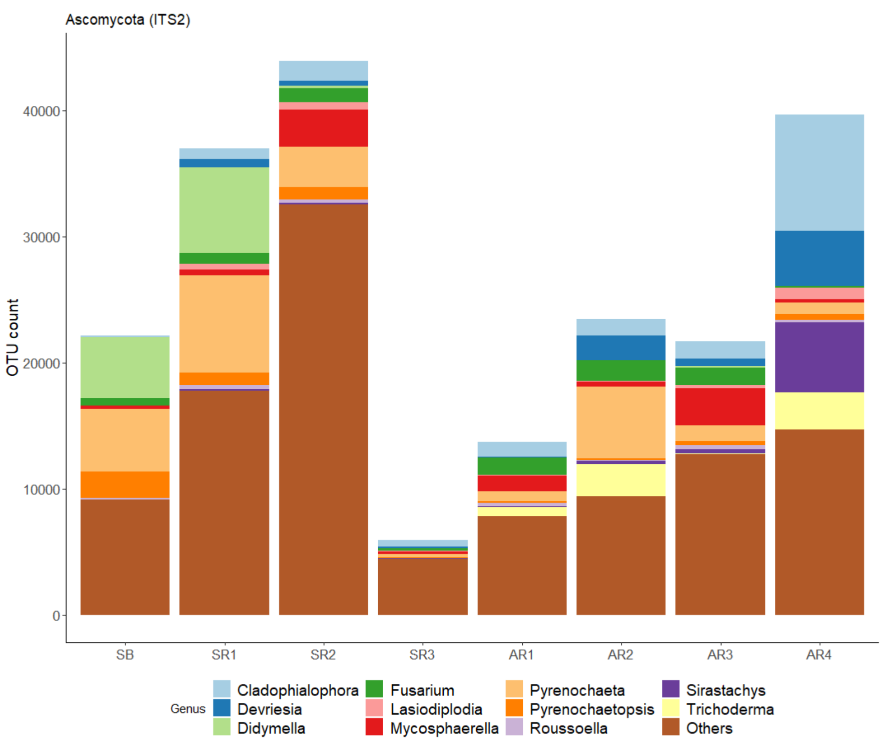


D

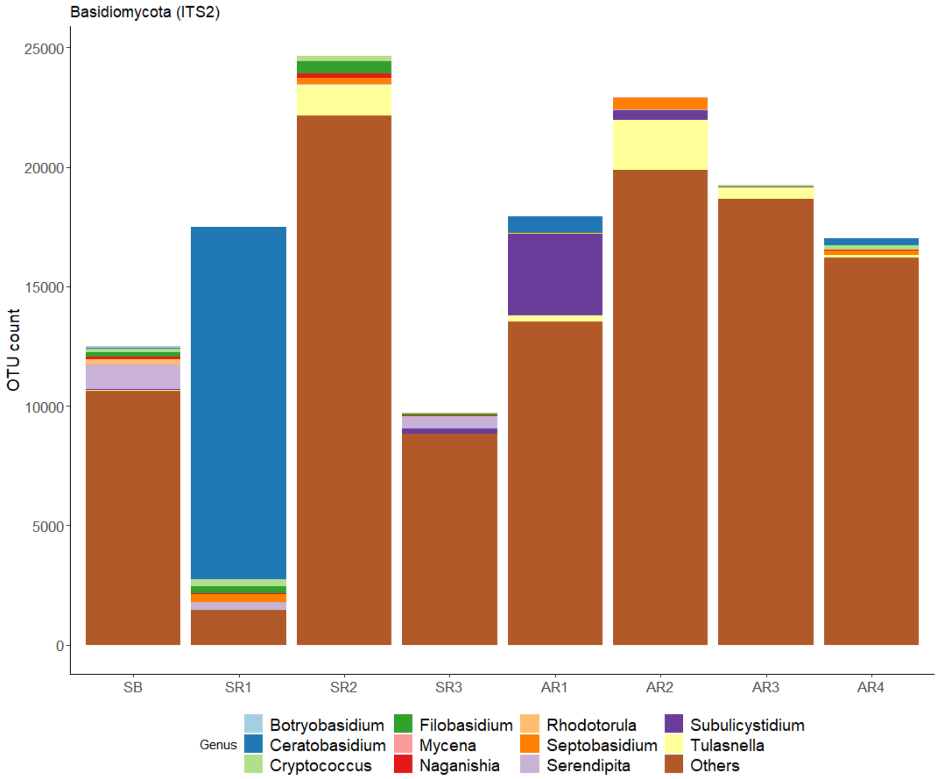


**Fig. S4 cont.**

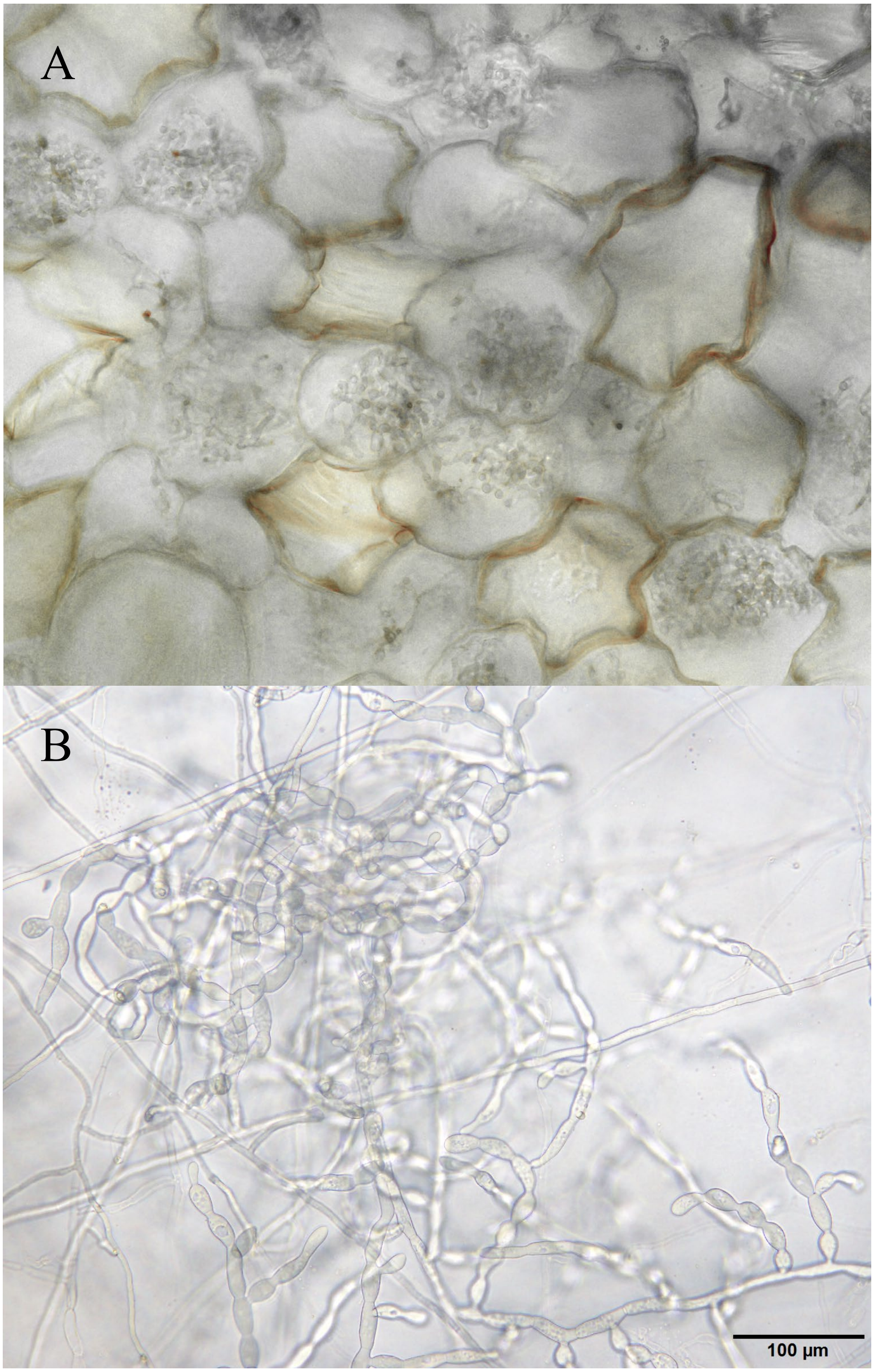


**Fig. S5.** A) Presence of *Rhizoctonia*-like fungus inside root cortex as intact pelotons with irregular hyphal width, resembling monilioid cells. B) Isolated *Rhizoctonia*-like fungi with conspicuous monilioid cells.

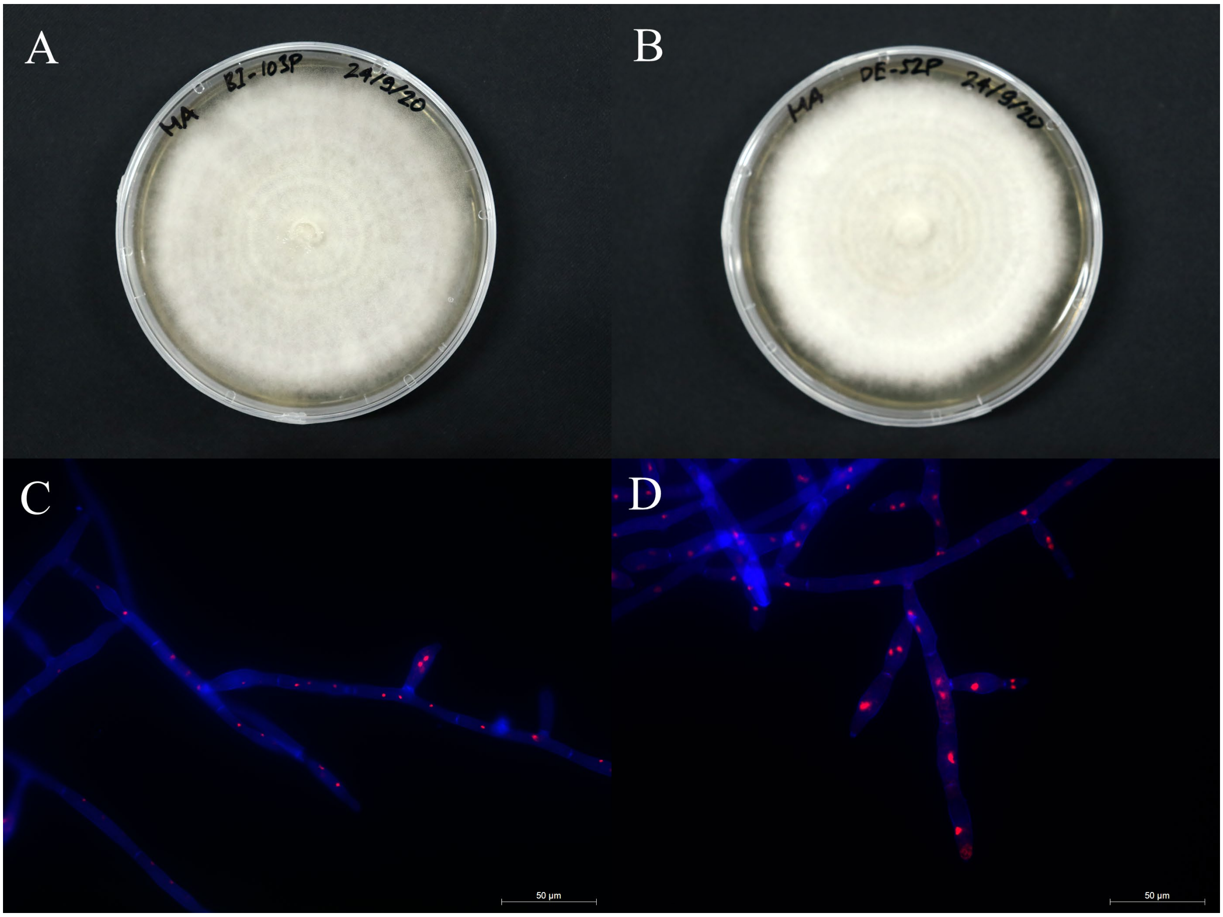


**Fig. S6.** Isolated *Rhizoctonia*-like fungi from roots. A , C) BI-103P and DE-52P cultures in malt extract agar (MA) after 2 weeks. B, D) Binucleate hyphae of BI-103P and DE-52P with branching at right angle.
